# A Robust Backscatter Modulation Scheme for Uninterrupted Ultrasonic Powering and Back-Communication of Deep Implants

**DOI:** 10.1101/2022.08.23.503752

**Authors:** Lukas Holzapfel, Vasiliki Giagka

## Abstract

Traditionally, implants are powered by batteries, which have to be recharged by an inductive power link. In the recent years, ultrasonic power links are being investigated, promising more available power for deeply implanted miniaturized devices. These implants often need to transfer back information. For ultrasonically powered implants, this is usually achieved with On-Off Keying based on backscatter modulation, or active driving of a secondary transducer. In this paper, we propose to superimpose subcarriers, effectively leveraging Frequency-Shift Keying, which increases the robustness of the link against interference and fading. It also allows for simultaneous powering and communication, and inherently provides the possibility of frequency domain multiplexing for implant networks. The modulation scheme can be implemented in miniaturized application specific integrated circuits, field programmable gate arrays, and microcontrollers. We have validated this modulation scheme in a water tank during continuous ultrasound and movement. We achieved symbol rates of up to 104 kBd, and were able to transfer data through 20 cm of water, with additional misalignment and during movements. This approach could provide a robust uplink for miniaturized implants that are located deep inside the body and need continuous ultrasonic powering.

## 1 Introduction

Conventional active implantable devices are powered by batteries that, when empty, need to either be replaced, or wirelessly recharged, usually inductively. Besides these conventional implants, a new family of wirelessly-powered, battery-free, active microimplants has emerged ([1], [2]). These aim to treat complex neurological disorders by providing access to various, and at times multiple, anatomical sites which can be located several cm deep inside the body. For this scenario, ultrasonic power transfer is being investigated due to its efficiency and miniaturization advantages ([3], [4], [5], [6], [7], [8]).

Active implants often need to communicate information, for example the power status of the implant or sensor data like neural signals [8] or blood pressure [9]. In conventional battery-powered active implants, Bluetooth Low Energy (BLE) is sometimes used for communication. This provides high data rates, at the expense of comparably high power consumption. Communication then might need a large portion of the total available power. For example, in [10] the authors needed a 90 mA h battery to achieve a discharge time of up to about 12 h, with most of the power being spent for the Bluetooth system-on-chip (SoC). This translates to a power consumption of several mW. Such battery sizes and power consumption ranges are often not acceptable for miniaturized implants. For power-aware or battery-less miniaturized implants, custom wireless electromagnetic or ultrasonic communication links are state-of-the-art. A comprehensive overview of existing methodologies for mm-sized networked implants can be found in [11], [12] and [13].

Electromagnetic links are widely used, however these are not suitable for miniaturized and deep im-plants. As explained in [11], the link frequency must be kept low to achieve low attenuation. But for low frequencies, in order to achieve sufficient coupling, the size of the coils must be at least the distance between the implanted and the external coils.

Generally, ultrasonic links have demonstrated higher penetration depths. We therefore turn to a fully-ultrasonic approach to power and communicate with deep implants. For a fully ultrasonic powerup and data uplink two approaches are generally used. A summary of these approaches is schematically illustrated in Figure 1 (left and middle). The first approach, known as backscatter modulation or load modulation, exploits the bidirectional characteristics of electro-acoustic transducers. First, the implant transducer converts the incoming acoustic power into electrical power. Modulating the load connected to the transducer alters the amount of electric power which is reflected back to the transducer. The transducer, in turn, converts this reflected electrical power back into acoustic power, effectively acting as a backscatterer. An external acquisition system can pick up this backscattered signal and detect the changes. A more detailed explanation of this concept can be found in [14].

**Figure 1:**
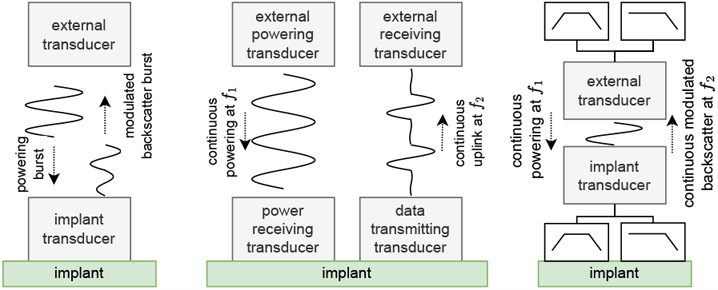
Available solutions for fully ultrasonic power-up and data uplinks. **left**) Backscatter modulation with time domain multiplexing. **middle**) Dedicated transducers with frequency multiplexing. **right**) Backscatter modulation with frequency domain multiplexing.

A second approach uses two distinct transducers, one dedicated to the power link, and an additional one for the communication uplink, as in [15] and [16]. The two links operate at different frequencies. The implant then incorporates an active circuit for driving the transducer used for communication. Such a system is described in more detail in [16].

Another approach was proposed recently in [17]. In this work, the authors combine backscattering with Frequency Division Multiplexing (FDM). With this approach, the implant transducer picks up a signal comprising two frequencies. One frequency is used for powering and another frequency for the uplink communication. The data uplink is subsequently separated from the power link via hardware filters at the implant.

For all approaches, variants of Amplitude Modulation (AM) [18], On-Off Keying (OOK) [16], or sometimes Pulse Width Modulation (PWM) [19] are used to encode the information.

The ultrasound backscatter modulation as a basis of OOK or AM (Fig. 1-left) typically requires pulsed ultrasound [18]. Also, determining the decision boundary between the high and the low level of the backscattered ultrasound can be difficult. Recent papers mitigate this problem by measuring the difference between two consecutive pulses ([4] and [19]), at the cost of dividing the data rate by two. On the other hand, adding a second, actively driven, transducer, as in Fig. 1-middle, increases the system complexity and size [18]. The same applies for the approach in [17], described in Fig. 1-right, as it requires at least two additional inductors and capacitors for the filter network. These limitations become more relevant for small, low-power implants that are placed deep inside the body.

To overcome these, we propose an approach based on combining backscattering with Frequency-Shift Keying (FSK). Compared to ASK or OOK, this increases the robustness of the communication link against noise [20]. We will show that FSK is also advantageous compared to ASK when considering interference and a fading channel, for example, due to patient movements. Similar approaches have already been described for inductive links ([21], [12]). A similar approach for ultrasonic uplinks was also recently proposed in [18] by Ghanbari, Piech, Shen, *et al*., but the subcarriers were used as a means of Code Division Multiplexing (CDM), to receive data from several im-plants simultaneously. Our approach would also allow for multi-device communication based on FDM or CDM, by choosing different modulation frequencies for each device. However, we only experimented with a single device. Importantly, our proposed approach allows for continuous ultrasound, using only a single transducer for power reception and communication on the implant side. Furthermore, it adds little complexity to the design of an application specific integrated circuit (ASIC) or field programmable gate arrays (FPGAs) and can also be implemented on microcontrollers.

## 2 Proposed modulation scheme and implementation considerations

### 2.1 Frequency-Shift Keying by load modulation (FSK-lm) uplink for ultrasonic biotelemetry

The proposed modulation scheme is based on the consideration that an implant needs to transfer back digital data and to be powered continuously, but the design requires a modulation scheme based on load or backscatter modulation, for example, due to size, complexity, or power constraints. In the presence of powering ultrasound, the detection of the backscattered AM signal becomes difficult, as its amplitude may be several orders of magnitude smaller than the incident ultrasound [18]. To mitigate this problem, we propose modulating the backscattered ultrasound at specific frequencies, to generate sidebands in the spectrum, which can be distinguished from the incident signal in the frequency domain. Fig. 3 shows the most straightforward implementation of this concept, which is superimposing a single subcarrier onto an OOK signal, shown in Fig. 2 b).

**Figure 2:**
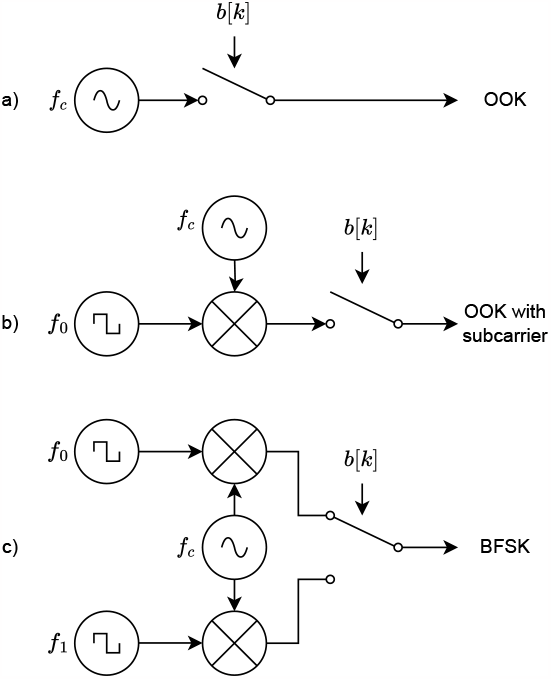
Signal flow diagrams of the mentioned modulation schemes. **a**) On-Off Keying with carrier frequency *f*_*c*_ **b**) On-Off Keying modulation with square subcarrier modulation frequency *f*_0_, used for example in near field communication (NFC), **c**) Binary Frequency-Shift Keying by switching between two different square subcarriers. The switch is toggled depending on the current bit *b*[*k*].

**Figure 3:**
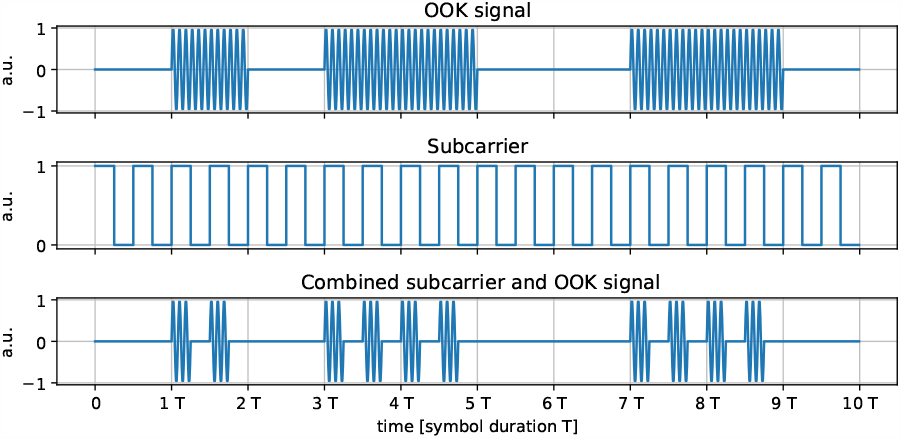
Simulated comparison of simple OOK and OOK with a superimposed subcarrier. For simple OOK the carrier amplitude is kept constant during each symbol duration. The subcarrier causes several short bursts with a constant repetition rate being generated during a single symbol.

Superimposing the subcarrier onto the OOK modulated signal simplifies the detection and decoding of the backscattered signal in the presence of a strong interference at the carrier frequency, as shown in Fig. 4. In the frequency domain, copies of the signal at frequencies further away from the incident ultrasound are present, providing the possibility to leverage FDM or CDM, as was done in conjunction with analog AM in [18]. It must be noted that the amplitude of these signal copies is inherently smaller compared to the amplitude of the simple OOK signal. Looking at Fig. 3, one can imagine the subcarrier effectively turning off the carrier signal half of the time. This means, on average, only half of the power is transferred when OOK is combined with a rectangular subcarrier compared to simple OOK. This already reduced power is spread across the carrier and the modulation products, causing further reduction of the amplitudes in the spectrum.

**Figure 4:**
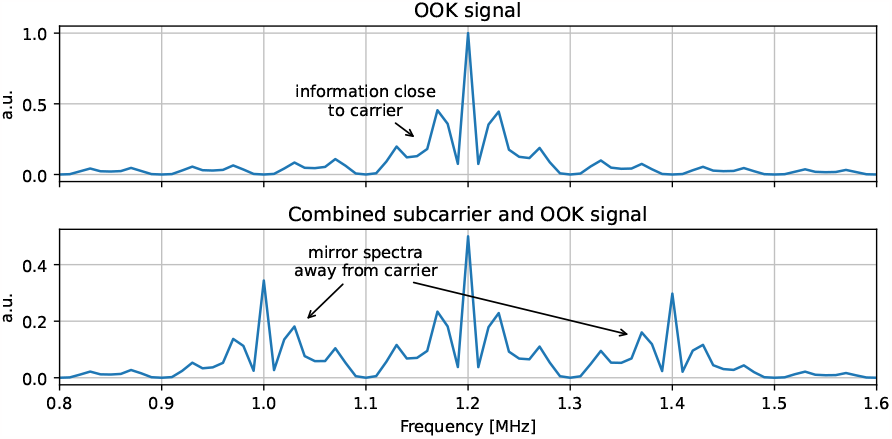
Simulated comparison of a simple OOK signal with a subcarrier superimposed OOK signal in the frequency domain. For simple OOK the information of the signal is spread in the frequency content close to the carrier. This makes it susceptible to fluctuations of the carrier, that alter the frequency content around it. Superimposing the OOK signal with a subcarrier generates mirrored signals further away from the carrier, which are therefore more robust against carrier fluctuations. Due to the additional modulation, the amplitude decreases when adding subcarrier modulation.

Combining OOK with a subcarrier would allow for continuous ultrasound powering and simultaneous communication. However, to further simplify the detection, we used several subcarriers that are selected depending on the data, effectively behaving as FSK. In the case of two distinct frequencies, we use one frequency *f*_0_ to encode a binary zero and another frequency *f*_1_ to encode a binary one, as shown in Fig. 2 c). This is equivalent to binary Frequency-Shift keying (BFSK). However, we do not actively drive the transducer with the two frequencies, but generate them through load modulation again. Therefore, we refer to it as Frequency-Shift Keying by load modulation (FSK-lm). Fig. 5 compares this modulation scheme with OOK in the time domain.

**Figure 5:**
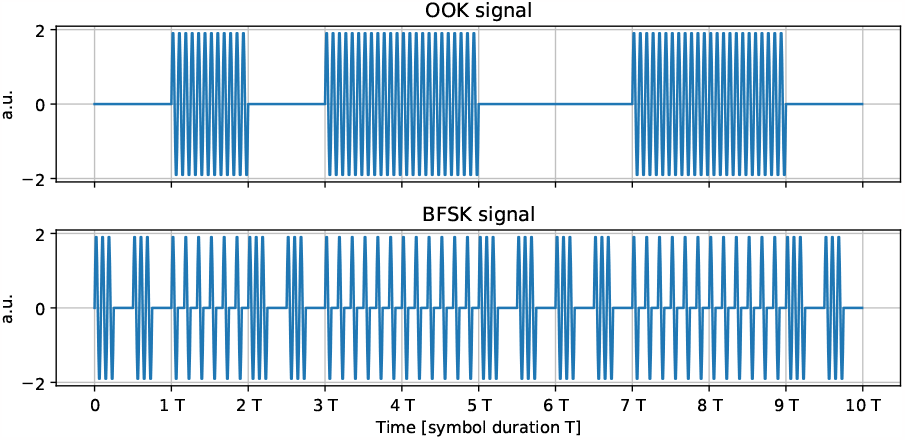
Simulated comparison of OOK and BFSK by load modulation in the time domain. For OOK, the carrier is turned on depending on the value of the current bit. For BFSK, the carrier is always pulsing, only the pulse rate is changed depending on the current bit value.

### 2.2 OOK versus BFSK with multiple backscatterers

Consider a more realistic channel as shown in Fig. 6 (top). Apart from the modulating transducer, another secondary backscattering surface is located nearby (e.g. bone tissue or the surface of the implant). The backscattered bursts from the modulating transducer and nearby surface may interfere. This can cause bit errors in case of OOK that can be avoided with BFSK (and FSK in general). In both cases, the secondary burst suddenly changes the amplitude of the received signal compared to the modulated signal from the implant. For OOK, this invalidates the decoding levels *A*_0_ and *A*_1_, causing a bit error. This could be mitigated, by e.g. sending a reference burst to remove the interference [19], or subtracting the constant interference during continuous power delivery. The above techniques can only be used if the channel is static, but this is unfortunately not the case within the human body [13]. In contrast, for BFSK such bit errors can be avoided even in non-static channels, as only the amplitudes of the superimposed modulation pulses with frequencies *f*_0_ and *f*_1_ are compared to each other, while the changing offset due to the secondary scattering is ignored. Of course, the effects of the secondary burst cannot be completely suppressed with FSK in case it occurs somewhere in the middle of a symbol, which, in reality, is always the case. It is, nevertheless, inherently suppressed by FSK demodulation, and the robustness against this phenomenon can be adjusted by the ratio of the modulation frequencies to the symbol rate. This property becomes more important for deep implants where the received modulated signal may be comparably weak to that of scatterers closer to the external transducer.

**Figure 6:**
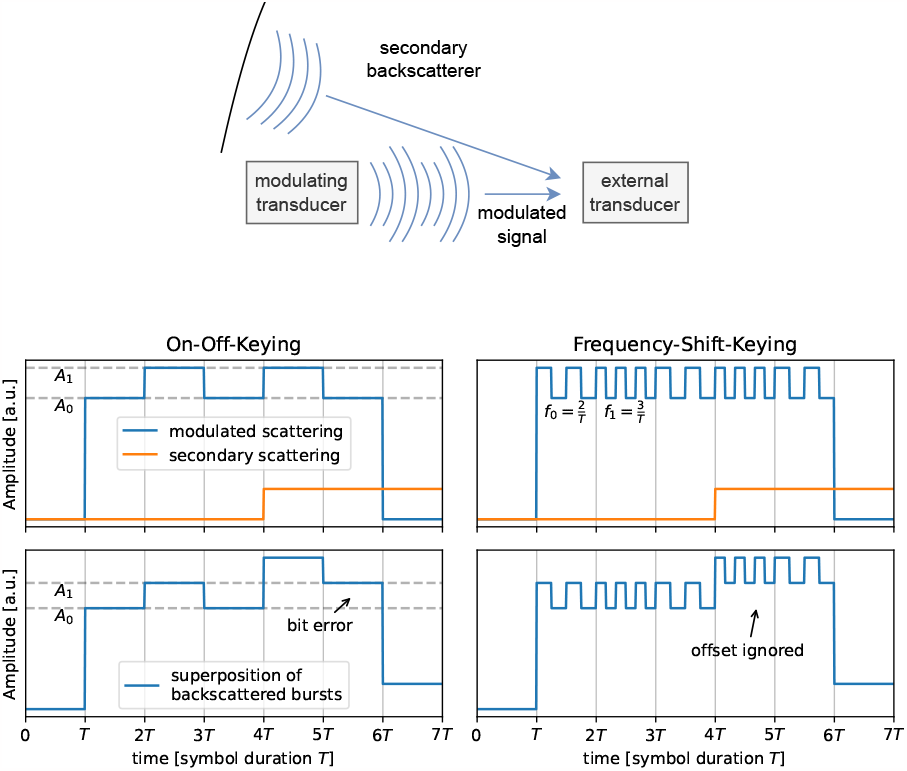
**top**) Model to compare ASK/PWM with FSK in the presence of a second backscattering surface. Depending on the distances of the modulating transducer and the secondary backscatterer to the external transducer, the backscattered bursts can interfere. **bottom**) Simplified simulation of the amplitude of backscattered signals for OOK and BFSK in the presence of a secondary scatterer.

### 2.3 BFSK signal generation in ASICs and microcontrollers

To generate the necessary control signal for the load switch, an ASIC or FPGA implementation would require one or more counters, depending on the specific modulation scheme (subcarrier, BFSK, or multiple Frequency-Shift Keying (MFSK)) and a few logic gates. Fig. 7 shows a simplified system overview needed for binary FSK.

**Figure 7:**
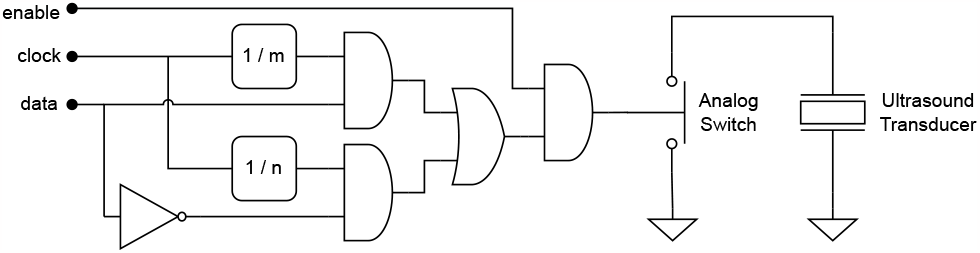
Simplified overview of the components needed to get a BFSK signal by load modulation. A clock signal is fed to two clock dividers, using integer denominators *m* and *n* with *m* ≠ *n*. The input signal “data” is a binary data stream. The logic gates select the modulation frequency depending on the bit value in the data stream. The enable signal is optional and might alternatively shut off the whole circuit, when no communication is needed.

Alternatively, a microcontroller-based implementation could be considered, as modern MCUs are able to generate the necessary control signal for the switch. They only need a timer peripheral that generates the control signal. The modulation frequency is then changed by updating the timer’s compare and period values.

## 3 Methods

To demonstrate the feasibility of this modulation scheme, we developed a bench top prototyping platform, which uses analog switches controlled by a timer peripheral of a microcontroller, to short circuit the transducer.

First we adapted the impedance matching network of the prototyping platform to the capacitive micromachined ultrasound transducer (CMUT), which was used as receiver on the implant side. We used one channel of a CMUT array described by Kawasaki, Westhoek, Subramaniam, *et al*. in [22]. The total area of one transducer channel is 6.3 mm^2^. As it is precharged, we did not need to apply a bias voltage. Fig. 8 shows the electrical impedance of the CMUT over frequency with the connected matching load.

**Figure 8:**
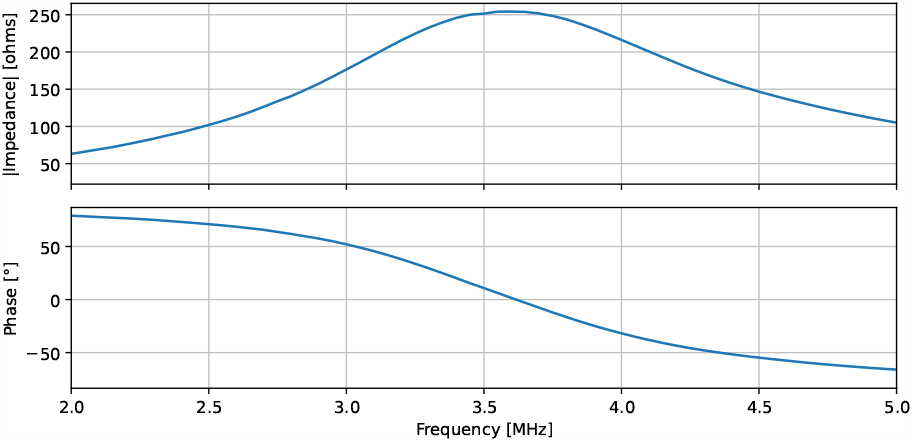
Measured electrical impedance of the CMUT, when connected to the matching network.

We then used this prototype in three different experiments. With the first, we investigated possible data rates and with the second we explored the out-of-focus performance. With the third experiment we transferred pseudo random data packages using both OOK and FSK, while moving the backscattering transducer, comparing the bit error ratio (BER).

All experiments were performed in a water tank and share the same setup, shown in Fig. 9. A commercial immersible TX transducer directs incident ultrasound with a frequency of 3.5 MHz to the CMUT, which is connected to the load modulation prototyping platform. The backscattered ultrasound is then picked up by a needle hydrophone. We also considered to use one transducer for both power transfer and data reception. However, with the given setup the dynamic range of the oscilloscope was not sufficient to detect the backscattered signal using a single transducer, with the data rates we anticipated. The prototyping platform and the setup are described in more detail in [23].

**Figure 9:**
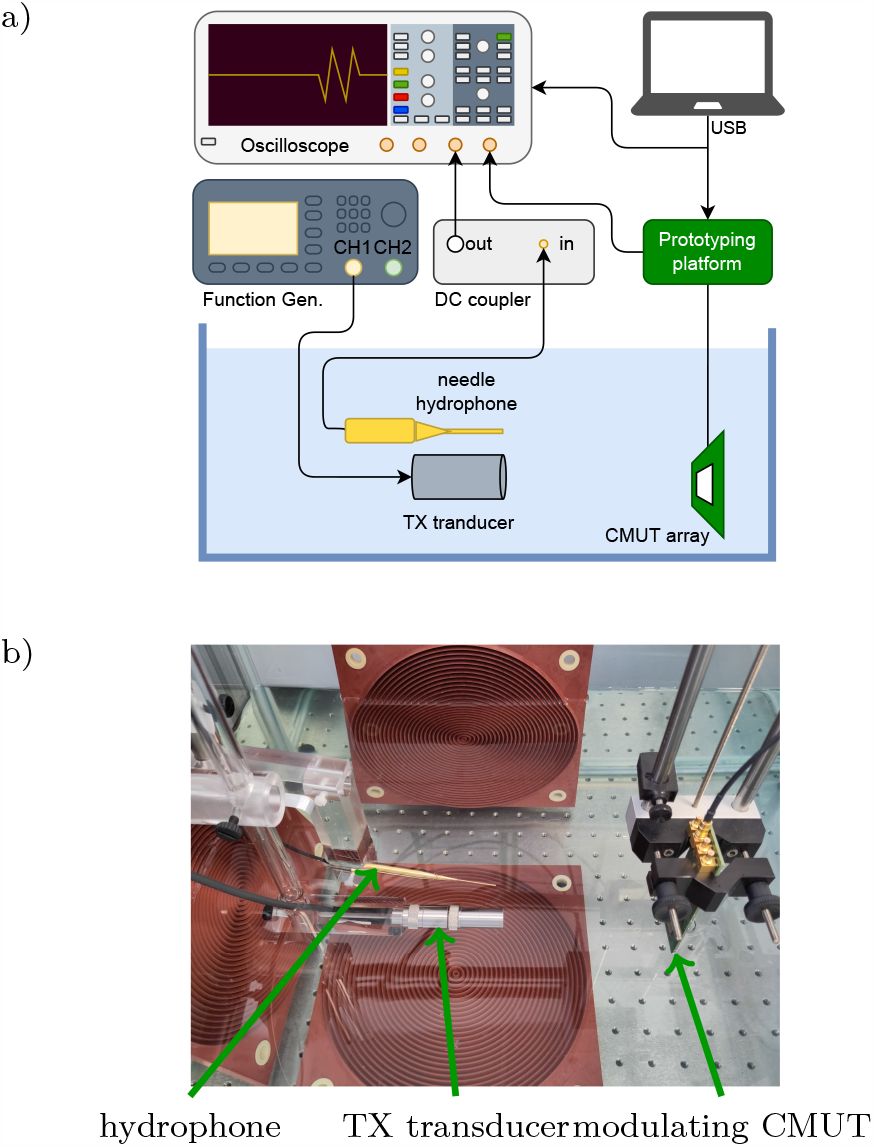
Setup of the performed experiments, from [23]. **a**) Schematic of the setup. The synchronization of the function generator and the prototyping platform to reference clock output of the oscilloscope is not shown. **b**) Arrangement of the transducers in the water tank.

For the transfer rate experiment, the CMUT was kept at the same position, but we increased the symbol rate until the system reached its limit. We also increased the encoding frequencies accordingly to at least two times the symbol rate.

For the out-of-focus experiment, we chose a fixed modulation frequency of 200 kHz and moved the CMUT to the side for 4 cm in 1 mm steps. How-ever, measuring the exact angles and distances of the setup was not possible. At each step, we recorded the hydrophone signal for 1.2 ms, corresponding to a symbol rate of 833 Bd, for offline analysis.

For the pseudo random data transfers we first improved the impedance matching. The resulting impedance is shown in Fig. 8. We then set the lateral start position of the CMUT to 0 mm or 5 mm away from the focal axis and the axial distance to approximately 12 cm or 20 cm. In total, this gives four different start positions. For each position the transfers were performed with a lateral velocity of 0 mm s^−1^, 1.8 mm s^−1^ and 18 mm s^−1^. The same transfer was repeated using OOK and FSK. For both encoding schemes we changed the symbol rate between 1.042 kBd and 104.17 kBd.

Using integer multiples of the symbol rate as modulation carrier frequencies, we made sure all frequencies are orthogonal, and do not leak into each other. This is especially important to suppress leakage from the strong interference at the carrier frequency into the relatively close-by frequency content of the modulated signal. For example, with a symbol rate of 104.17 kBd, the carrier frequency was set to *f*_*c*_ = 34 · 104.17 kHz = 3.54 MHz, and the modulation frequencies where set to *f*_0_ = 4 · 104.17 kHz = 416.67 kHz, and *f*_1_ = 6 · 104.17 kHz = 625 kHz respectively. We demodulated the signal based on the Goertzel algorithm. The receiver implementation is described in more detail in [23]. The packet size was chosen as a power of ten, depending on the symbol rate, to get a packet transfer duration close to 1 s. For example, for 1.042 kBd the packet size is chosen to be the closest power of ten, which is 10_3_ = 1000.

## 4 Results

### 4.1 Data transfer rate

For the data transfer rate experiment, we achieved symbol rates of 100 kBd, as shown in Fig. 10. With 150 kBd, the microcontroller was not able to update the switching frequency for the next symbol in time. The bandwidth and noise of the channel would have allowed for higher symbol rates.

**Figure 10:**
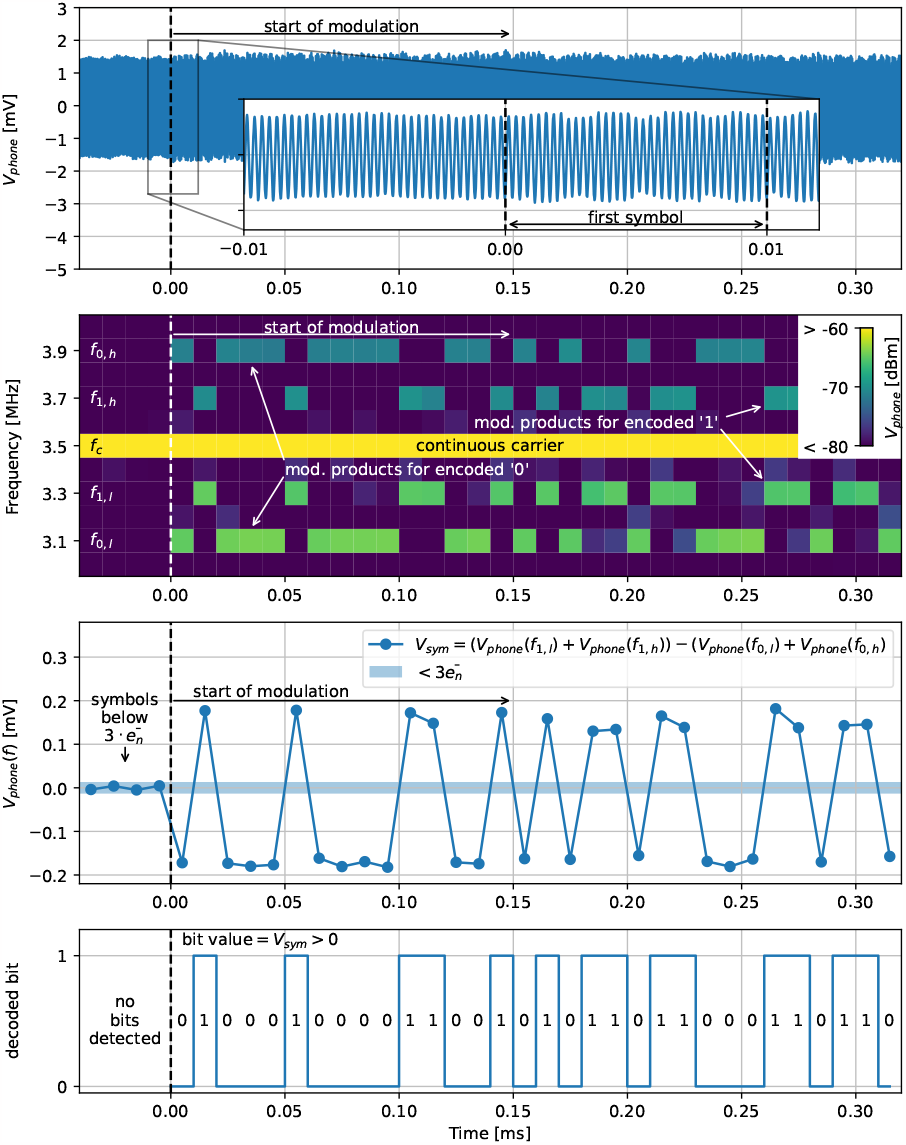
Decoding of a recorded FSK signal with a symbol rate of 100 kBd. Zeros are encoded with a modulation frequency of *f*_0_ = 400 kHz, ones with *f*_1_ = 200 kHz. **top**) captured hydrophone signal showing the beginning of the modulation at *t* = 0 ms. **middle top**) Spectrogram of the backscattered signal. From *t* = 0 ms on, the modulation products above and below the carrier frequency are visible. The signal range was compressed to visually enhance the modulation products at *f*_0,*l*_ = 3.1 MHz, *f*_1,*l*_ = 3.3 MHz, *f*_1,*h*_ = 3.7 MHz and *f*_0,*h*_ = 3.9 MHz. **middle bottom**) The bipolar signal calculated from the modulation products. **bottom**) The decoded binary data.

### 4.2 Out-of-focus performance

For the out-of-focus experiment, an initial 60 µs burst was sent, which allowed us to estimate the distances and how the ultrasound waves propagate with the CMUT on-axis. Fig. 11 shows the recorded signals of this burst.

**Figure 11:**
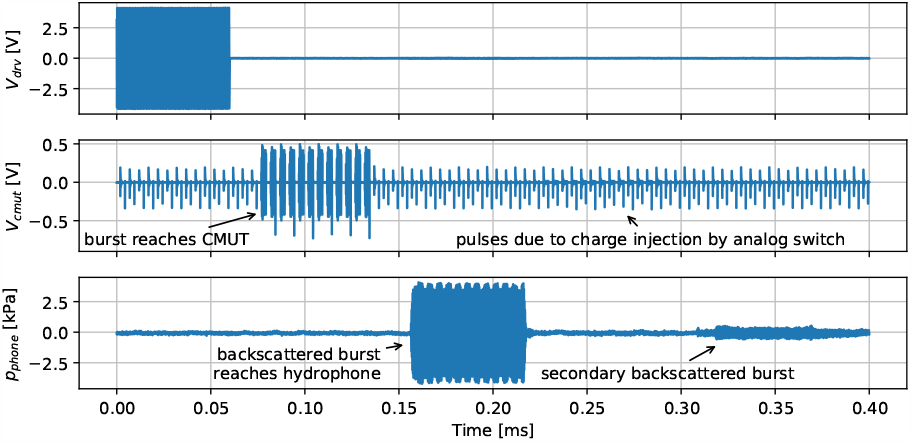
Burst to measure distances at the start position of the out-of-focus scan. **top**) The driving voltage of the external transducer reaches about 8 V_pp_. **mid**) After about 80 µs the burst reaches the CMUT. **bottom**) After approximately another 80 µs, the backscattered burst reaches the hydrophone with visible modulation. After about 0.32 ms, secondary backscatting arrives.

After about 80 µs, the burst reached the CMUT and after about another 80 µs, the hydrophone picked up the backscattered and modulated burst. For the voltage at the CMUT, one can see a periodic disturbance, also without the ultrasound burst being present. This is caused by the parasitic input capacitance of the analog switches MOSFETs, which results in brief current spikes flowing through the CMUT whenever the analog switches state is changed. To ensure that we did not accidentally misinterpret the ultrasound pulses generated by this disturbance as our modulation signal, we chose the modulation frequency such that the frequencies of the modulation products would not be harmonics of the modulation frequency itself. With a carrier frequency of *f*_*c*_ = 3.5 MHz, the closest modulation products become *f*_*c*_ − *f*_*mod*_ = 3.3 MHz and *f*_*c*_ + *f*_*mod*_ = 3.7 MHz. As these frequencies are not integral multiples of the modulation frequency *f*_*mod*_, we can be sure that we did not detect a harmonic of these pulses.

Then we moved the CMUT in 1 mm steps laterally out of the focal axis of the TX transducer. We calculated the amplitude of the modulation products and the carrier from the hydrophone signal for each step using an FFT. Fig. 12 shows the calculated amplitudes at the CMUT and the hydrophone depending on the position of the CMUT.

**Figure 12:**
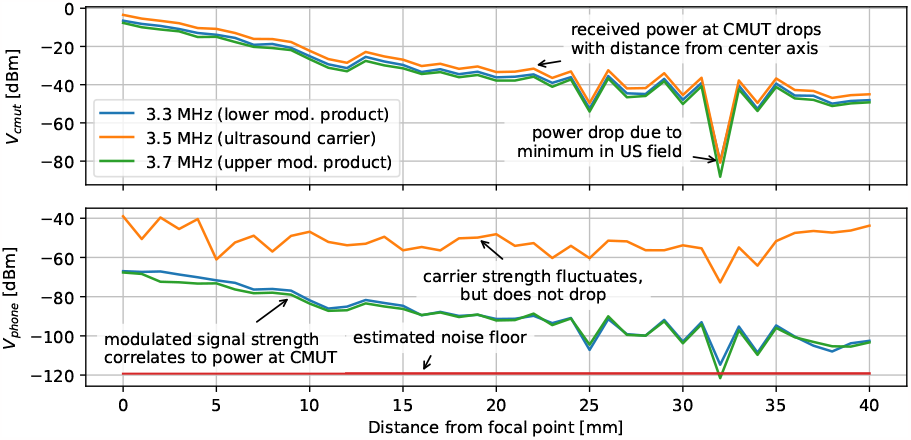
Measured signal strengths depending on the CMUT transducer’s distance to the TX transducer’s focal axis. **top**) Voltage across the CMUT transducer that modulates the ultrasound. As the modulation switch is shorting the CMUT, the modulation depth is close to 100 %, and the amplitude of the modulation products is close to the carrier amplitude. **bottom**) The backscattered signal, picked up by the hydrophone. While the amplitude of the modulation products has a visible correlation to their amplitude at the CMUT transducer, the carrier amplitude is mostly dependent on the backscattering of the surrounding setup.

The figure shows, that in our setup the amplitude of the modulation products remains 10 dB above the noise floor, with the CMUT array up to 4 cm away from the optimum position. The only exception is at 32 mm, where the received acoustic power at the CMUT drops by several orders of magnitude. We estimated the noise floor from the measured amplitude at frequencies away from the carrier and the modulation products.

### 4.3 Measurement of BER during movement

The results of the pseudo random data transfer experiment are summarized in Table 1. From the table it can be seen immediately that the combination of a symbol rate of 104 kBd and a velocity of 18 mm s^−1^ is not feasible for both modulation schemes (last two columns, first four rows in the table). Out of the remaining 31 scenarios, the BER for OOK is lower than for FSK, only for the position furthest away. This is due to additive white Gaussian noise (AWGN) dominating the adverse effects. To improve the BER the symbol rate can be reduced for FSK. In contrast, for OOK, reducing the symbol rate worsens the performance when movement is involved. When moving with 18 mm s^−1^ with OOK no usable data rate can be found.

**Table 1:**
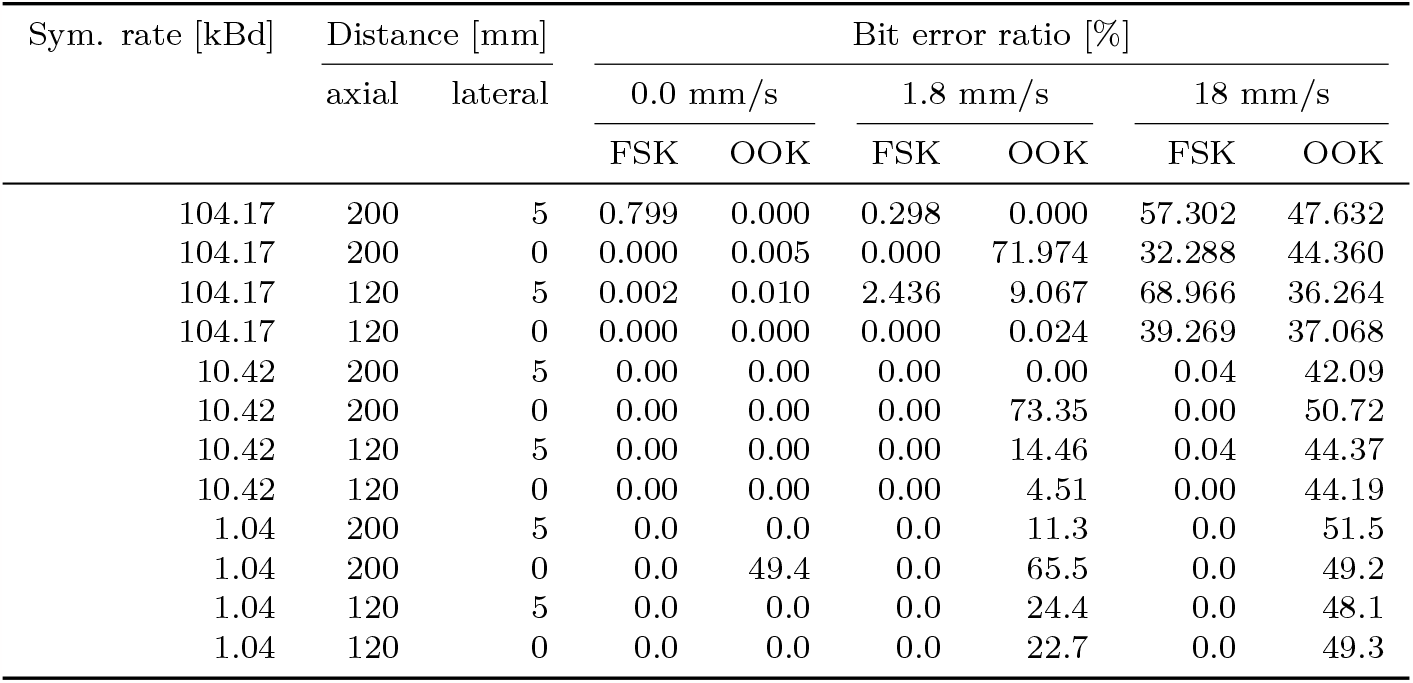
Comparison of the measured bit error ratio of OOK and FSK for different scenarios.

| Sym. rate [kBd] | Distance [mm] |  | Bit error ratio [%] |  |  |  |  |  |
| --- | --- | --- | --- | --- | --- | --- | --- | --- |
|  | axial | lateral | 0.0 mm/s |  | 1.8 mm/s |  | 18 mm/s |  |
|  |  |  | FSK | OOK | FSK | OOK | FSK | OOK |
| 104.17 | 200 | 5 | 0.799 | 0.000 | 0.298 | 0.000 | 57.302 | 47.632 |
| 104.17 | 200 | 0 | 0.000 | 0.005 | 0.000 | 71.974 | 32.288 | 44.360 |
| 104.17 | 120 | 5 | 0.002 | 0.010 | 2.436 | 9.067 | 68.966 | 36.264 |
| 104.17 | 120 | 0 | 0.000 | 0.000 | 0.000 | 0.024 | 39.269 | 37.068 |
| 10.42 | 200 | 5 | 0.00 | 0.00 | 0.00 | 0.00 | 0.04 | 42.09 |
| 10.42 | 200 | 0 | 0.00 | 0.00 | 0.00 | 73.35 | 0.00 | 50.72 |
| 10.42 | 120 | 5 | 0.00 | 0.00 | 0.00 | 14.46 | 0.04 | 44.37 |
| 10.42 | 120 | 0 | 0.00 | 0.00 | 0.00 | 4.51 | 0.00 | 44.19 |
| 1.04 | 200 | 5 | 0.0 | 0.0 | 0.0 | 11.3 | 0.0 | 51.5 |
| 1.04 | 200 | 0 | 0.0 | 49.4 | 0.0 | 65.5 | 0.0 | 49.2 |
| 1.04 | 120 | 5 | 0.0 | 0.0 | 0.0 | 24.4 | 0.0 | 48.1 |
| 1.04 | 120 | 0 | 0.0 | 0.0 | 0.0 | 22.7 | 0.0 | 49.3 |

For the scenarios with an axial distance of 20 cm and a lateral distance of 0 mm, the BER of OOK is exceptionally high. Without movement and with the lowest symbol rate of 1.04 kBd, the BER for OOK reaches almost 50 %, in this case. Near that position, the combination of the phase and amplitude differences between the useful signal and the interference “hides” the amplitude modulation of the useful signal at the receiver. This effect is so sensitive to the CMUT position, that the precision of the positioning causes it to happen only in one out of three scenarios (with same nominal position but different symbol rates). With added movement of 1.8 mm s^−1^ the BER for OOK increases well above 50 % for all symbol rates. In those scenarios, the movement brings the transducer into a region, where destructive interference between the secondary scattering and the useful signal occurs, effectively inverting the signal at the receiver. We described above effects on OOK in more detail in [23].

## 5 Discussion

In this study, we have proposed Frequency-Shift Keying by load modulation for ultrasonic uplink communication from small and deep implants. Our proposed approach, as opposed to time-multiplexed schemes as in Fig. 1-left, allows ultrasonic communication with uninterrupted power transfer, thus maximizing the amount of power that can reach the implant. In contrast to [18], the additional subcarrier is not used to differentiate between several implants, but instead to differentiate the backscattered signal from the incident ultrasound. Our approach would also allow for multi-device communication based on FDM, by choosing different frequencies for each device. A similar modulation approach, using a single subcarrier, is standardized for inductive Near-Field Communication (NFC) of identification/proximity cards in ISO/IEC 14443-2.

The frequency of the powering ultrasound, depends on the geometry of the ultrasound link and the transducers in use. Thus, finding the optimum modulation frequencies for FSK and symbol rate is a challenge, that has to be uniquely solved for each system or use case. With this scheme, the subcarrier frequency could be adapted on the fly.

The experimental validation of this scheme comes with some practical limitations, which are discussed below.

The power transfer efficiency of our setup is not maximized due to the TX transducer not being used at its resonant frequency. Pre-charged RX CMUTs can achieve efficiencies of 50 % [24]. Assuming random data, on average, the power consumption of FSK is close to OOK, only the additional power consumed by the circuit needed to generate the FSK signal as shown in Fig. 7 has to be added. This would be only a fraction of the mW-range instantaneous power which was recently demonstrated ultrasonically [25]. However, for OOK the efficiency depends on the data itself, whereas for FSK, the switch is closed 50 % of the time at most, regardless of the data.

The proposed communication scheme has been validated using a benchtop setup, and practical choices related to this validation, such as the equipment used, will unavoidably affect and often determine the measured experimental performance. For example, in this study we have demonstrated symbol rates of up to 104 kBd. This symbol rate can be limited by the bandwidth of the transducer or the impedance matching network. In general FSK-lm modulation needs a higher bandwidth than OOK for the same symbol rate. To achieve orthogonal modulation frequencies with non-coherent detection, the modulation frequencies must be integer multiples of the symbol rate. For BFSK this means the necessary bandwidth is increased at least by a factor of 2.

In this implementation, the main limiting factor for the symbol rate was the time the microcontroller needed to update its timer peripheral for the next symbol. Increasing its core clock frequency, using a faster microcontroller, or leveraging direct memory access (DMA) techniques would have allowed for higher data rates. Other studies with a focus on millimeter scale implants have reported data rates of 40 kbit s^−1^ ([11] and [16]), 95 kbit s^−1^ with dedicated transducers for the data uplink [15], and a possible 200 kbit s^−1^ in [17], derived from the pulse response of the system. In [4], the symbol rate is in the order of 1 kBd, but depends mainly on the pulse repetition rate. If size and implant depth are of no concern, ultrasound communication is capable of achieving data rates in the range of 100 Mbit s^−1^ ([26] and [27]). We expect our scheme to also be capable of reaching several Mbit s^−1^ in an experimental setup, if optimized for maximum data rate, ignoring other constraints, such as power, size, complexity, depth, etc. How-ever, in this case, using other modulation schemes could be more advantageous as they would allow for even higher data rates and better robustness, as has been done in [26] and [27]. Overall, for similar scenarios comparable rates to the literature are achievable with our approach.

However, the focus of our approach is not high data rates, but the relatively high robustness combined with minimum added complexity and versatility. To demonstrate this robustness, we have considered scenarios where out-of-focus detection is required and movements are involved. We were not able to find any relevant reports on the out-of-focus performance or involved movements in the literature. We assume the reason for that is that perfect alignment is necessary/implied for systems benchmarked so far. Comparing the robustness of our scheme to that achieved by OOK/ASK modulation, we can see that the main advantage of our approach can be derived from the ratio between the received modulation amplitude and the fluctuations of the carrier amplitude. This ratio can reach several orders of magnitude. For OOK, the high ratio makes it difficult for the decoding algorithms to let the decision boundary track the fluctuations. Also destructive interference between the secondary scattering and the useful signal can cause high bit error ratios for OOK, even in a static channel. This issues is completely avoided, when using FSK.

It should also be noted, that, as long as the implant load is not changing with frequencies close to the modulation frequency, the only effect of the implant on the communication link will be a changing modulation index resulting in a varying signal strength. The same is true for OOK, where the load could be changing with the symbol rate. However, FSK has the advantage, that the modulation frequencies can be chosen -to some degree- independently from the symbol rate. So FSK may be used to mitigate problems that arise from load fluctuations of the implant itself.

As an additional remark, it should be noted that all experiments were performed in a water tank, i.e. in a very controlled environment. In a real world scenario, the medium would be tissue instead of water, which has higher attenuation and scatters ultrasound itself. We could also expect larger misalignment and scattering from other surfaces or implants. This could reduce the SNR, which means the distances or data rates achieved in this experiment might not be feasible. However, the outlined advantages over OOK should apply in just these scenarios. Finally, in the current implementation, all signals were recorded using an oscilloscope and analyzed offline later. Future work will have to show online decoding to reduce saved data size and allow for closed loop systems.

In addition, the performance of other modulation schemes like MFSK, Phase-Shift Keying (PSK), or Quadrature Amplitude Modulation (QAM) can possibly be implemented with our prototyping platform and may improve performance. For example in [20], binary Phase-Shift Keying (BPSK) is shown to have a lower BER than BFSK. Adding Manchester coding could simplify the clock synchronization. The effects of multi path propagation can be further investigated. A future version of the prototyping platform could be powered from the ultrasound carrier itself, to show the feasibility of combining both links.

## 6 Conclusion

In this work, we have proposed a data communication scheme based on Frequency-Shift Keying by load modulation (FSK-lm), to enable robust ultrasonic uplink communication during continuous ultrasonic power transfer in small and deep implants. A single transducer on the implant side is sufficient for simultaneous power reception and communication. We were able to transfer data through water with distances up to 20 cm, with additional misalignment and during movements. Our experiments showed that FSK-lm, compared to the more common OOK/ASK approaches in ultrasonic uplinks, increases the communication robustness, while achieving similar symbol rates. We used an off-the-shelf microcontroller to generate the necessary control signal for the load modulation switch and showed that integration into an ASIC requires only a few additional circuits.

## Acknowledgement

This work is part of the Moore4Medical project funded by the ECSEL Joint Undertaking under grant number H2020-ECSEL-2019-IA-876190.

